# Analysis of correlation and trail coefficients for componentperformance s in nine experimental tomato lines

**DOI:** 10.1101/2021.03.18.436039

**Authors:** Gonzalo Quispe Choque

## Abstract

The objective of the research was to analyze the main variables related to tomato yield, and guide the selection of materials for the INIAF vegetable improvement program. The experiment was carried out in the open field using nine tomato lines on the grounds of the National Vegetable Seed Production Center, during the 2017-2018 agricultural campaign. A randomized complete block experimental design was used, with three repetitions and 10 plants per experimental unit. For the analysis of the data, the variable yield was considered as dependent and the variables number of flowers per inflorescence, number of clusters per plant, number of fruits per plant, weight of fruit, equatorial and polar diameter as independent variables. Analysis of variance, phenotypic correlations and path coefficients were performed. The performance of the L015 line was 80. 79 t ha-1 higher than the L014, L019 and Rio Grande lines. The fruit yield had a significant correlation with the weight of fruit per plant followed by the polar diameter, equatorial diameter, number of fruits per plant and weight of fruit. The analysis of path coefficients showed that the number of fruits per plant had the highest direct positive effect on the fruit yield, fruit weight and equatorial diameter that have a significant correlation and a direct effect on the fruit yield, emerged as the components with coefficients of 0.96 and 0.52 respectively. These characters may be relevant within the selection criteria in the development of new varieties. number of fruits per plant and weight of fruit. The analysis of path coefficients showed that the number of fruits per plant had the highest direct positive effect on the fruit yield, fruit weight and equatorial diameter that have a significant correlation and a direct effect on the fruit yield, emerged as the components with coefficients of 0.96 and 0.52 respectively. These characters may be relevant within the selection criteria in the development of new varieties. number of fruits per plant and weight of fruit. The analysis of path coefficients showed that the number of fruits per plant had the highest direct positive effect on the fruit yield, fruit weight and equatorial diameter that have a significant correlation and a direct effect on the fruit yield, emerged as the components with coefficients of 0.96 and 0.52 respectively. These characters may be relevant within the selection criteria in the development of new varieties.

## Introduction

Tomato (2n = 24), belonging to the Solanaceae family, is a important vegetable in the world with a potential of Performance 33.98 t ha^-1^ and a kind well studied in terms of genetics (Foolad, 2007 and FAOSTAT, 2018). This fact derives from the different types of fruits that the species presents and the varied forms of consumption that it offers., particularly like a rich source vegetable of carotenoidsvitamins, carbohydrates, as well as other essential minerals (Bergougnoux, 2014; Schwarz et al., 2014; Giovannucci et al., 2002 and Patiño et al., 2015). In Bolivia, its cultivated areaa in the main producing regions It is 4691 hectares with a production from 61,360 to 63,454 t year^-1^ and a yield per unit area of 12 to 13 t ha^-1^ that is less than the half its potential performance. These data demonstrate the greatimportance socioeconomic of this crop (OAP, 2019).

Systematic study and evaluation of germplasm tomato is of great importance for agronomic and genetic improvement current and future cultivation (Reddy et al., 2013). In general, when working with tomato cultivation, a large number of variables are measured to obtain a data set that allows the most varied types of statistical evaluations and analyzes. When numerous variables are studied at the same time, correlations between them can be calculated, which are important for the selection of characteristics of interest for plant breeding (Moreira et al., 2013). Without embargo, the acquaintancethe relationship between andl performance and other characters of the plant and its relative contribution to performance it is very useful when formulating the selection scheme. As performance is a complete characterjor, it is difficult to explore multiple characters that contribute to the same to through of the correlation, so so much, it is important to carry out other analyzes that include the coefficients de path that provide a clear indication for the selection criteria. In this way, the path analysis is a statistical analysis capable of recognizing cause and effect relationships (Wright, 1921), displaying the correlation coefficients in the direct and indirect effects of the independent variables in a dependent variable.

National Vegetable Project of the National Institute of Agricultural and Forestry Innovation, comes identifying high-yielding genotypesin fruit, quality and tolerance to adverse abiotic and biotic factors, evaluating a large number of variables that allow to explain performance components by medium of a simple model, analyzing its numerical components, such as the number of fruits per plant that is determined by the number of flowers that are fertilized and the final weight of the same. This work aimed to analyze the main variables related to tomato yield, and guide the selection of materials for the INIAF vegetable improvement program.

## Materials and methods

Essay It has been made in the National Center for Vegetable Seed Production of the INIAF, locatedat municipality of Sipe Sipe, Quillacollo province, of the Cochabamba department. Geographically it is located 17°26’24.4” South latitude; 66 ° 20’38.9” west longitude and at a height of 2505 m.s.n.m, during the 2018-2019 agricultural season.

For the development from work sand they used seven experimental tomato lines, to this material was addedor two varietyit is As a Witness (Lia and Rio Large), in order to compare the superiority or inferiority of the materials in terms of productivity (Table 1). The sowing of the genetic material was carried out in multicell trays of 128 alveothe under glass, with rice husk, lama and topsoil as substrate. LThe seedlings were transplanted 36 days after sowing in open field conditions when they presented cinco true leaves, using a stocking density of 20,000 pl ha^-1^.Lthe plants were tutored when they reached 15 cm Tall, the leaf removal of the lower leaves was carried out once the fruits of the first cluster were formed. I knowused drip irrigation 20 cm apart, with two daily irrigations of 20 min each, applying approximately 1.13 l per plant day 1. The fertilizers were applied by fertigation with direct suction through a Venturi, the daily doses were according to the phenological stage of the crop, the total applied was: 260N-330P-330K. The fruit harvest began 75 days after transplantation, manually, once a week.

**Table 1.** Origin and agronomic characteristics of lines experimental tomato tested during the 2018-2019 agricultural season at the INIAF National Vegetable Seed Production Center, Cochabamba, Bolivia.

| No. | Experimental Line | Origin | Fruit | Cycle |
| --- | --- | --- | --- | --- |
| 1 | L014 | PNH (INIAF) | Oblong | Early |
| 2 | L015 | PNH (INIAF) | Ovoid | Semi early |
| 3 | L027 | PNH (INIAF) | Round | Semi early |
| 4 | L031 | PNH (INIAF) | Round | Semi early |
| 5 | L019 | PNH (INIAF) | Oblong | Semi early |
| 6 | AVTO1003 | AVRDC (Taiwan) | Oblong | Semi early |
| 7 | AVTO1007 | AVRDC (Taiwan) | Square | Semi early |
| **Cultivars** |  |  |  |  |
| 8 | Rio Grande | CNPSH (INIAF) | Piriform | Early |
| 9 | Lia (tester) | Sakata | Piriform | Early |

PNH: National Vegetable Project. INIAF, Cochabamba, Bolivia
AVRDC: Asian Vegetable Research and Development Center. Shanhua, Taiwan.
CNPSH: National Center for Vegetable Seed Production. INIAF, Cochabamba, Bolivia

I know usedA statistical design of complete random blocks, with nine treatments (experimental lines) and three repetitions, the experimental unit consisted of 10 plants distributed in 2 rows, 80 cm apart and 2 m long each. For harvest purposes, 5 plants were taken per experimental unit. Variables associated with fruit yield components of the second cluster were evaluated in five individual plants per plot, in free competition. Table 2 describes the name of the response variables, symbol, and units of measurement; These were evaluated according to the descriptor manual for tomato (S. lycopersicum L.) ofl International Plant Genetic Resources Institute (IPGRI, 1996) and the guide of the International Union for the Protection of New Variety of Plants for tomato (UPOV, 2011).

**Table 2.** Fruit yield response variables and their components of the genotypes tested during the 2018-2019 agricultural season at the INIAF National Center for Vegetable Seed Production, Cochabamba, Bolivia.

| No. | Variable | Symbol | Measurement units | Measuring instrument |
| --- | --- | --- | --- | --- |
| 1 | Number of flowers per inflorescence | NFI | unite | counting |
| 2 | Cluster number per plant | NFR | unite | counting |
| 3 | Number of fruits per bunch | NFR | unite | counting |
| 4 | Equatorial diameter | OF | mm | vernier |
| 5 | Polar diameter | DP | mm | vernier |
| 6 | Fruit weight | Pf | g | precision scale |
| 7 | Number of fruits per plant | NFP | unite | counting |
| 8 | Fruit weight per plant | PFP | kg | precision scale |
| 9 | performance | RDTO | t ha<sup>-1</sup> | precision scale |

The performance and its components were analyzed by vari analysisanza, considering genotypes as fixed effects. When the significance levels were p≤ 0.05 averages were calculated and the test of averages of Minimum Significant Difference (LSD). The phenotypic correlations (r) between the variables were calculated using Pearson’s correlation coefficient, and the path coefficient analysis was performed, considering fruit yield as a dependent variable and yield components as independent variables. All the analyzes were carried out with the softwaresstatistics: SPSS V23 (2014) R Project 3.2.5. (2016).

## Results and Discussion

The mean square values of the variance analysis for characters fruit weight (PF), fruit equatorial diameter (DE), polar diameter (DP), fruit weight per plant (PFP), number of fruits per plant (NFP) and yield (RDTO) subjected to study, revealed highly significant differences (p≤0.01) and for the number of bunches per plant (NRP) and number of fruits per bunch (NFR) significant differences (p≤0.05), between the lines studied (Table 2). These results indicate that the differences are due to the intrinsic genetic conditions of each cultivar. The experimental variation coefficients due to their low value (<27%) reveals the existence of experimental precision, which allows guaranteeing the validity of the conclusions reached. Similar results were reported by Dar and Sharma (2011); Jilani et al. (2013); Monamodi et al. (2013).

Table 4 presents the results of the mean test. The highest number of fruits per bunch was the lines AVTO1007, AVTO1003, L014 and Rio Grande. While L015, Lia, L027 and L029, showed the lowest amounts. For the variable average fruit weight, line L015presented the highest weight and lastly L027 with the best fruit weight. González and Laguna, (2004) cited by Blandon (2017) affirm that the differences in fruit weight between the genotypes are due to the genetic makeup of each line and the influence exerted by the environment. The most outstanding lines in performance were L015,Bundle, L14 and Rio Grande. The fact that these lines developed a greater number of fruit per plant and with a greater fruit weight stands out. Ponce (1995), indicates that the number of fruits per plant is associated with their morphological parts; thus, the number depends to a great extent on the type of inflorescences possessed by the cultivars, be they simple or compound.

**Table 3.** Analysis of variance (ANVA) for nine quantitative characters evaluated in nine experimental tomato lines during the 2017-2018 agricultural season.

| sv | DF | Mean Square |  |  |  |  |  |  |  |  |
| --- | --- | --- | --- | --- | --- | --- | --- | --- | --- | --- |
|  |  | NFI | NRP | NFR | PF | DE | DP | PFP | NPF | RDTO |
| **Blq** | 2 | 0.78 * | 0.70 ** | 0.44<sup>ns</sup> | 271.47 * | 1.46<sup>ns</sup> | 1.18<sup>ns</sup> | 0.21<sup>ns</sup> | 110.70 * | 57.07<sup>ns</sup> |
| **Line** | 8 | 0.75<sup>ns</sup> | 0.28 * | 1.58 * | 1856.87 ** | 46.48 ** | 70.73 ** | 4.12 ** | 448.54 ** | 969.68 ** |
| **E. Exp.** | 16 | 0.94 | 0.25 | 0.78 | 205.80 | 6.32 | 4.46 | 0.97 | 101.37 | 143.33 |
| **CV%** |  | 15.90 | 13.37 | 15.56 | 18.43 | 5.08 | 3.43 | 26.68 | 20.67 | 20.17 |

Number of fruits per inflorescence (NFI), Number of clusters per plant (NRP), Number of fruits per cluster (NFR), Fruit weights (PF), equatorial diameter of fruit (DE), Polar diameter of fruit (DP), Fruit weight per plant (PFP), Number of fruits per plant (NPF), Fruit yield per plant (RDTO).
\* Significant at $p < 0.05$ ; \*\* Significant at $p < 0.01$ ; ns not significant

**Table 4.**
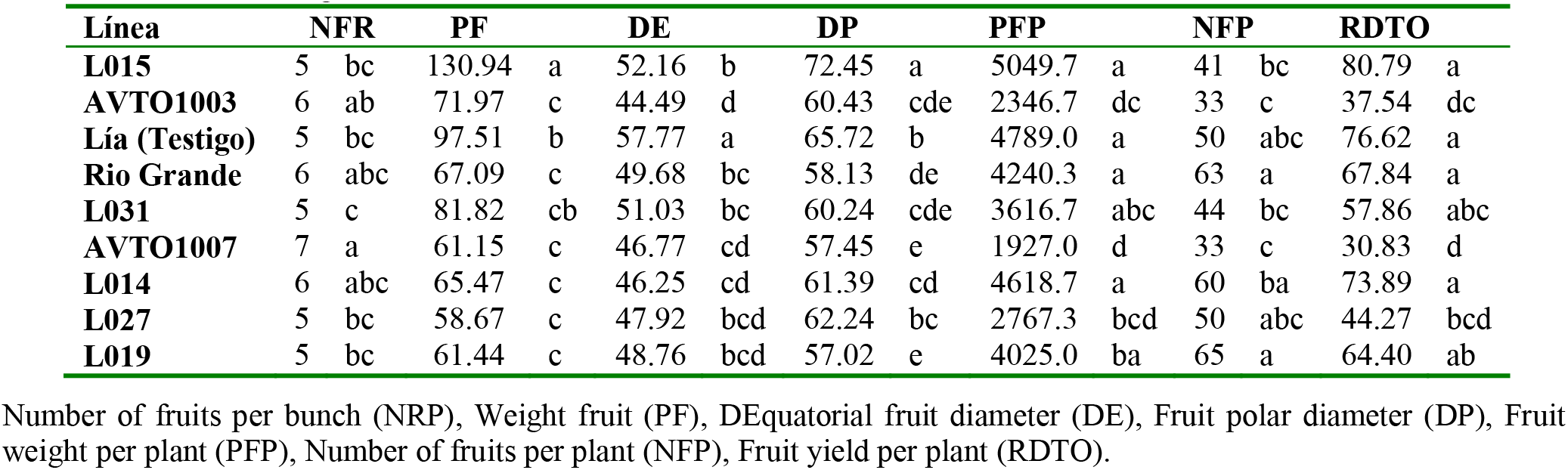
Comparison of means of LSD for nine quantitative characters evaluated in nine experimental tomato lines during the 2017-2018 agricultural season.

| Línea | NFR |  | PF |  | DE |  | DP |  | PFP |  | NFP |  | RDTO |  |
| --- | --- | --- | --- | --- | --- | --- | --- | --- | --- | --- | --- | --- | --- | --- |
| **L015** | 5 | bc | 130.94 | a | 52.16 | b | 72.45 | a | 5049.7 | a | 41 | bc | 80.79 | a |
| **AVTO1003** | 6 | ab | 71.97 | c | 44.49 | d | 60.43 | cde | 2346.7 | dc | 33 | c | 37.54 | dc |
| **Lía (Testigo)** | 5 | bc | 97.51 | b | 57.77 | a | 65.72 | b | 4789.0 | a | 50 | abc | 76.62 | a |
| **Rio Grande** | 6 | abc | 67.09 | c | 49.68 | bc | 58.13 | de | 4240.3 | a | 63 | a | 67.84 | a |
| **L031** | 5 | c | 81.82 | cb | 51.03 | bc | 60.24 | cde | 3616.7 | abc | 44 | bc | 57.86 | abc |
| **AVTO1007** | 7 | a | 61.15 | c | 46.77 | cd | 57.45 | e | 1927.0 | d | 33 | c | 30.83 | d |
| **L014** | 6 | abc | 65.47 | c | 46.25 | cd | 61.39 | cd | 4618.7 | a | 60 | ba | 73.89 | a |
| **L027** | 5 | bc | 58.67 | c | 47.92 | bcd | 62.24 | bc | 2767.3 | bcd | 50 | abc | 44.27 | bcd |
| **L019** | 5 | bc | 61.44 | c | 48.76 | bcd | 57.02 | e | 4025.0 | ba | 65 | a | 64.40 | ab |

Number of fruits per bunch (NFR), Weight fruit (PF), DEquatorial fruit diameter (DE), Fruit polar diameter (DP), Fruit weight per plant (PFP), Number of fruits per plant (NFP), Fruit yield per plant (RDTO).

The phenotypic correlations between the variables were calculated using Pearson’s correlation coefficient. Such correlations constitute a measure of the magnitude of the linear association between two variables without considering cause and effect between them regardless of the units. The relationship between traits is generally due to the presence of linkages and the pleiotropic effect of different genes. In Figure 1, The values of the simple correlations obtained between pairs of variables are shown. Among the variables of yield components, the highest correlation corresponded to the weight of the fruit with the polar diameter of the fruit (r= 0.80**), these variables, in their order, are highly correlated with fruit weight per plant and fruit length (r= 0.68** and r= 053**). Positive correlations of the number of flowers per inflorescence with the number of fruits percluster (r= 0.92 **), and equatorial diameter (r= -0.42 **) indicate that plants tend to develop more of these characteristics causing smaller diameter fruits. Kaushik and Singh (2018) reported similar results. Without embargo, Escalante (1989), indicatesthat the larger the fruit, the less the number of fruits. This is corroborated by the characteristics of each cultivar since the photosynthates that the plant assimilates in some cases increase the number of fruits and in others increase the size. Antonio andSolis (1999), showed that when the weight of the fruit increased, the number of them per plant was reduced, with a negative correlation.

**Figure 1.**
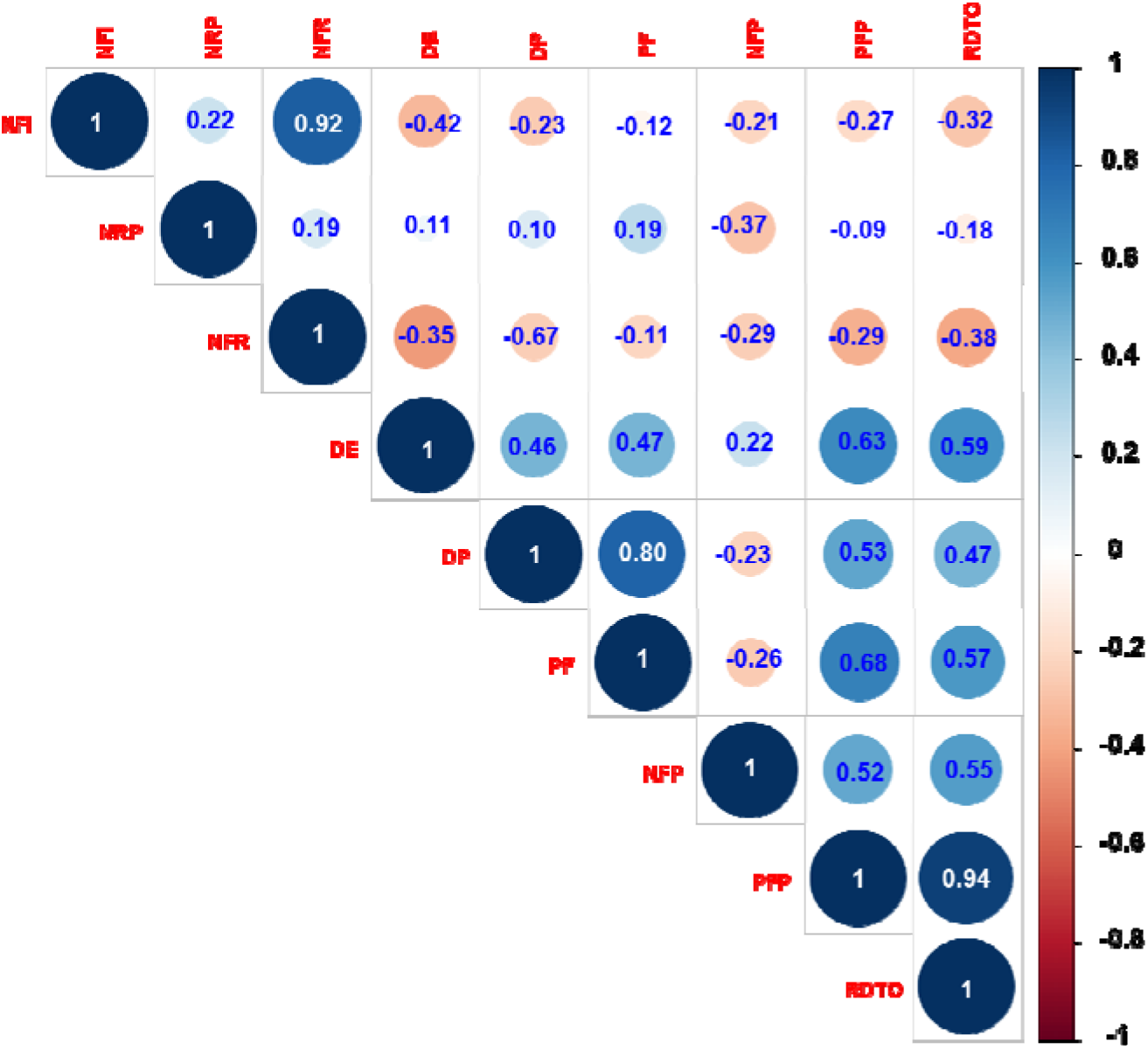
Correlogram of the degree of association between fruit yield and its components in experimental tomato lines, evaluated during the 2018-2019 agricultural season.

Among the variables that are highly correlated with fruit yield are fruit weight per plant (r= 0.93**),average fruit weight (r= 55 **), equatorial diameter of the fruit (r= 0.63 **) and polar diameter of the fruit (r= 0.47*). Indicating that the lines with greater development in equatorial and polar diameter tend to have higher fruit weights. Cancino (1990) found that fruit size (closely related to fruit weight) depends on three to five pairs of genes, an aspect that agrees with what Ashcroft et al. (1993), in which the size of the fruit is controlled by genetic factors, in addition to physiological factors; such as ripening, topping and defoliation.

Tiwari and Upadhyay (2011) reported that the height of the plant, the diameter of the fruit and the length of the fruit were directly responsible for determining the fruit yield in tomato. Haydar et al(2007) also observed that fruit weight exerted a high positive and direct effect on fruit yield per plant

Trail analysis is a reliable statistical technique, devised by Wright (1921), that helps determine the traits that contribute to performance and is therefore useful in indirect selection. Provides possible explanations for the correlations observed between a dependent variable and a series of independent variables, separating the direct effects of one variable on another and the indirect effects of one variable on another via one or more independent variables and helps the breeder to determine the components performance.

The coefficients of the path analysis in Figure 2 indicated that the fruit weight, number of fruits per plant and the diameter equatorial had a maximum direct contribution of (P_PF RDTO_ = 0.52, P_NFP RDTO_ = 0.96 and P_DP RDTO_ = 0.52) together with a highly significant correlation with fruit yield. Traits showing a high direct effect on yield per plant indicated that direct selection could be effective in improving yield based on selection of these traits. In this regard, Singh and Chaudhary (1985) cited by Duarte et al., (2012), indicate that being positive (both direct effects and correlation coefficients), the correlation explains the true relationship between these characters and a direct selection to through these characteristics it will be effective.Similar results obtained Monamodi et al. (2013),who when evaluating six lines of tomato with a determined habit found that the yield per plant was positively correlated with the number of fruits per bunch (r = 0.59), number of bunches per plant (r = 0.87), number of fruits per plant (r = 0.90), fruit weight per bunch (r = 0.59). De Souza et al., (2012) also found that the fruit yield per plant was positively related to the variables number of fruits per plant (r = 0.94), average fruit weight (r = 0.53), number of clusters per plant (r = 0.72) and number of fruits per bunch (r = 0.82).

**Figure 2.**
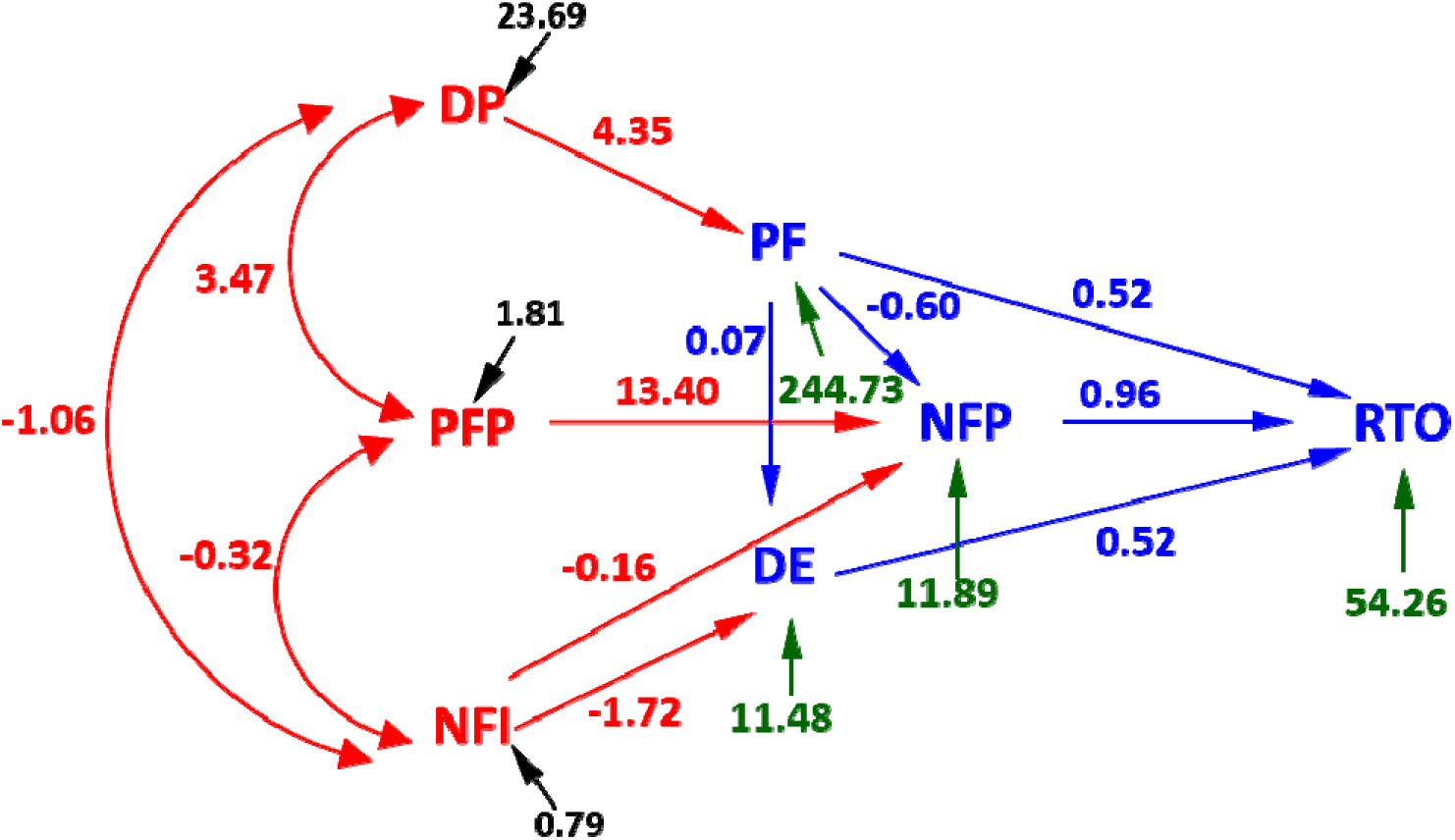
Path analysis diagram indicating the direct and indirect effects of the yield components on the yield of the experimental tomato lines, evaluated during the 2018-2019 agricultural season.

He effect indirect number of flowers per inflorescence by via diameter equatorial and number of fruits per plant it was of *P*_NFI DAND_= -1.72 and P_NFI NFP_=-1.16 respectively, indicating that few genetic gains can be achieved in the selection process for plants of greater number of flowers per inflorescence, due to its low contribution to fruit yield. Followed by the indirect effect via number of fruits per plantand diameter equatorial with coefficients of -0.60 and -0.07 and finally the equatorial diameter has an indirect effect on the yield by fruit weight (4.35).

## Conclusion

It was found that the yield it isvariable between tomato lines. The presence of this variability is important because the success of any crop improvement depends on the variability and, to a greater extent, on the parameter that is heritable. The experimental lineL015, emerged as a materialbetter in terms of performance and most of the components measured compared to the control. Fruit weight is associated with the variables polar diameter, equatorial diameter and fruit weight per plant, with which high phenotypic correlation values were obtained. In the analysis of trail coefficients, it revealed that the number of fruits per plant had a greater direct effect on yield. Fruit weight was the second most important component with a better direct effect. Therefore, it can be concluded that the aforementioned characters should be duly considered when formulating the selection strategy to develop high-yielding tomato varieties.

## Notes

### Competing Interest Statement

The authors have declared no competing interest.

